# Acute myeloid leukemia stratifies as two clinically relevant sphingolipidomic subtypes

**DOI:** 10.1101/2023.04.13.536805

**Authors:** B. Bishal Paudel, Su-Fern Tan, Todd E. Fox, Johnson Ung, Jeremy Shaw, Wendy Dunton, Irene Lee, Arati Sharma, Aaron D. Viny, Brian M. Barth, Martin S. Tallman, Myles Cabot, Francine E. Garrett-Bakelman, Ross L. Levine, Mark Kester, David Claxton, David J. Feith, Kevin A. Janes, Thomas P. Loughran

## Abstract

Acute myeloid leukemia (AML) is an aggressive disease with complex and heterogeneous biology. Although several genomic classifications have been proposed, there is a growing interest in going beyond genomics to stratify AML. In this study, we profile the sphingolipid family of bioactive molecules in 213 primary AML samples and 30 common human AML cell lines. Using an integrative approach, we identify two distinct sphingolipid subtypes in AML characterized by a reciprocal abundance of hexosylceramide (Hex) and sphingomyelin (SM) species. The two Hex-SM clusters organize diverse samples more robustly than known AML driver mutations and are coupled to latent transcriptional states. Using transcriptomic data, we develop a machine-learning classifier to infer the Hex-SM status of AML cases in TCGA and BeatAML clinical repositories. The analyses show that the sphingolipid subtype with deficient Hex and abundant SM is enriched for leukemic stemness transcriptional programs and comprises an unappreciated high-risk subgroup with poor clinical outcomes. Our sphingolipid-focused examination of AML identifies patients least likely to benefit from standard of care and raises the possibility that sphingolipidomic interventions could switch the subtype of AML patients who otherwise lack targetable alternatives.

**Key Points:** 1.Sphingolipidomics separates acute myeloid leukemia (AML) patients and cell lines into two subtypes.

2.The subtype with low hexosylceramide and high sphingomyelin defines a new high-risk subtype with poor clinical outcomes.

## Introduction

Recent work has combined proteomics^1^, signaling^2,3^, or immunophenotypes^4^ with integrated genomic-transcriptomic measurements to improve acute myeloid leukemia (AML) patient risk classifications beyond mutations and cytogenetics^5^. Although invaluable as resources, such approaches cannot extend retroactively to existing repositories nor prospectively to new AML cases lacking these data types. We sought to develop a more-extensible approach involving sphingolipids (**Figure 1A**), a family of bioactive molecules implicated in AML pathogenesis and therapeutic resistance^6,7^. Sphingolipid species are delicately balanced and several differentially regulate cell proliferation^8^, differentiation^9^, autophagy^10^, apoptosis^11^, and immune cell activation^12^. Recent evidence indicates that sphingolipid abundances in AML are heterogeneous^13^, prompting us to ask whether systematic sphingolipidomic profiling could meaningfully stratify AML patients and common AML cell lines.

**Figure 1:**
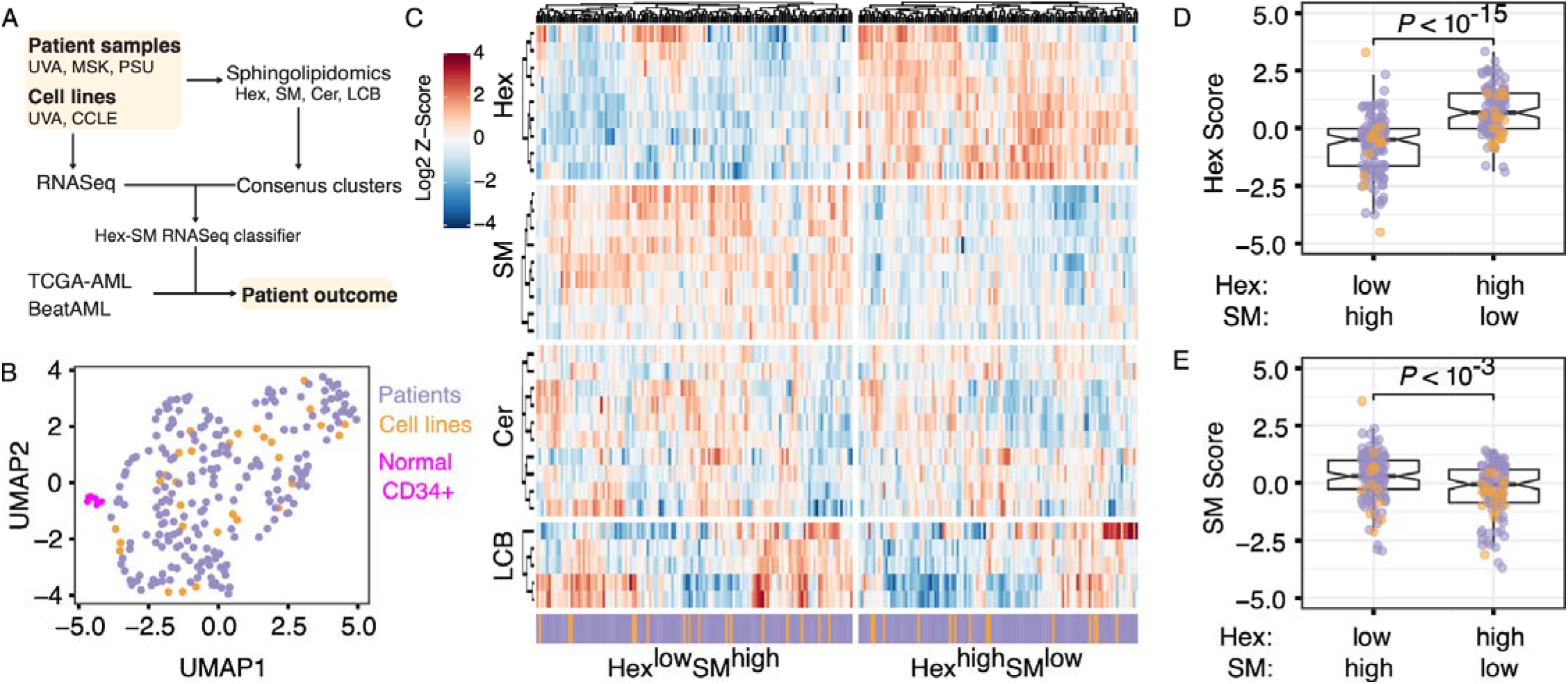
AML cell lines and patients separate into two sphingolipidomic clusters that differ in their abundance of hexosylceramide and sphingomyelin. (**A**) Strategy to identify sphingolipidomic subtypes in AML. Sphingolipidomics of ceramide (Cer), hexosylceramide (Hex), sphingomyelin (SM), and long-chain bases (LCB, comprised of sphingosine and its derivatives) was performed on primary AML samples and AML cell lines by LC-MS, and the normalized data were consensus clustered to identify a stable number of sphingolipid clusters. Cluster-specific gene signatures were extracted to train a Hex-SM classifier that infers sphingolipidomic subtype from RNASeq. (**B**) Sphingolipidomic heterogeneity is similar in AML cell lines and patient samples but distinct from normal CD34+ bone marrow. Normalized sphingolipidomics for normal bone marrow samples (magenta, *n* = 6), primary AML samples (purple, *n* = 213), and AML cell lines (orange, *n* = 30) were displayed by Uniform Manifold Approximation and Projection (UMAP). (**C**) Row-standardized lipid abundances organized by sphingolipid family: hexosylceramide (Hex), sphingomyelin (SM), ceramide (Cer), and long-chain bases (LCB). The Hex^low^SM^high^ and Hex^high^SM^low^ consensus clusters are separately clustered and annotated as cell lines (orange) and patient samples (purple). (**D-E**) Normalized Z-scores of lipid species within the Hex (**D**) and SM (**E**) families were summed and differences between consensus clusters were assessed by the Mann-Whitney test with continuity correction. Colors indicate the sample type: AML cell lines (orange, *n* = 30) and primary samples (purple, *n* = 213).

### Study Design

Patient samples were obtained from the University of Virginia Cancer Center (UVA), Memorial Sloan Kettering Cancer Center (MSK), and Penn State Hershey Cancer Center (PSU, **Figure 1A**). Targeted sphingolipidomics by mass spectrometry and transcriptomics by RNASeq were performed on both primary AML samples and cell lines. Transcriptomic data for TCGA-AML and BeatAML were downloaded from the National Cancer Institute Genomic Data Commons data portal. The Hex-SM classifier was developed as a support vector machine with a linear kernel and 60-40 training-test data allocation. More details on the Study Design and Methods are described in the Supplemental Information.

## Results and Discussion

We quantified 33 sphingolipid metabolites in 213 primary AML samples, 30 human AML cell lines, and 6 normal CD34+ enriched bone marrow samples after carefully controlling for cell purity and viability. Normalized sphingolipid profiles in AML cell lines and primary AML cases were highly dispersed (yet intermixed) and separable from normal samples (**Figure 1B**), motivating a pan-AML stratification. We applied consensus clustering to the normalized lipidomics data and identified two sphingolipidomic clusters that were statistically robust (**Supplemental Figure S1A-C**). The two clusters were equally populated with cell lines and primary samples, and neither was differentially enriched for common AML mutations (**Supplemental Figure S1D**, Supplemental Table 01; Fisher’s exact test, *P*_adj_ ≥ 0.27). In contrast, the clusters were divergent in their abundance of hexosylceramide (Hex) and sphingomyelin (SM) species (**Figure 1C**). Lipid cluster 1 exhibited proportionally less Hex and more SM (Hex^low^SM^high^), whereas cluster 2 exhibited more Hex and less SM (Hex^high^SM^low^, **Figure 1D-E)**. Additionally, the Hex^high^SM^low^ cluster was elevated in long-chain, C14-20-carbon chains relative to the Hex^low^SM^high^ cluster (**Supplemental Figure S1E**; Mann-Whitney test, *P* < 10^-9^). No differences were detectable in the abundance of ceramide (Cer), other long-chain/sphingoid bases (LCB, comprised of sphingosine and its derivatives), or very-long-chain, C22-26 chain sphingolipids (**Supplemental Figure S1F-H**; Mann-Whitney test, *P≥*0.20). Collectively, the analyses supported two sphingolipidomic subtypes with biochemical states that were largely uncoupled from the AML derivation or mutation status.

We next examined whether AML patients from the two subtypes differed in their clinical outcomes. Complete data across the three centers were available for 70 Hex^low^SM^high^ and 73 Hex^high^SM^low^ cases sampled at diagnosis before intensive induction chemotherapy (Supplemental Table 02). Based on European LeukemiaNet (ELN) 2022 criteria^5^, patients in the Hex^low^SM^high^ subtype had a twofold higher rate of failure compared to patients that were Hex^high^SM^low^ (**Supplementary Figure S2A**; Fisher’s exact test, *P* = 0.02). The Hex^low^SM^high^ subtype also trended towards shorter event-free survival (EFS) and overall survival (OS), although the difference was not statistically significant (**Figure 2A-B;** median EFS = 139 vs. 239 days, log-rank *P* = 0.16; median OS = 454 vs. 786 days, log-rank *P* = 0.38). To eliminate confounding caused by events that occurred before the first round of induction therapy was complete, we excluded patients with EFS < 20 days and observed significant differences in EFS and OS (**Supplemental Figure S2B-C;** median EFS = 142 vs. 339 days, log-rank *P* = 0.014; median OS = 454 vs. 1577 days, log-rank *P* = 0.03). These results were robust to the choice of EFS threshold (**Supplemental Figure S2D**) and suggested that the sphingolipidomic subtypes have prognostic value.

**Figure 2:**
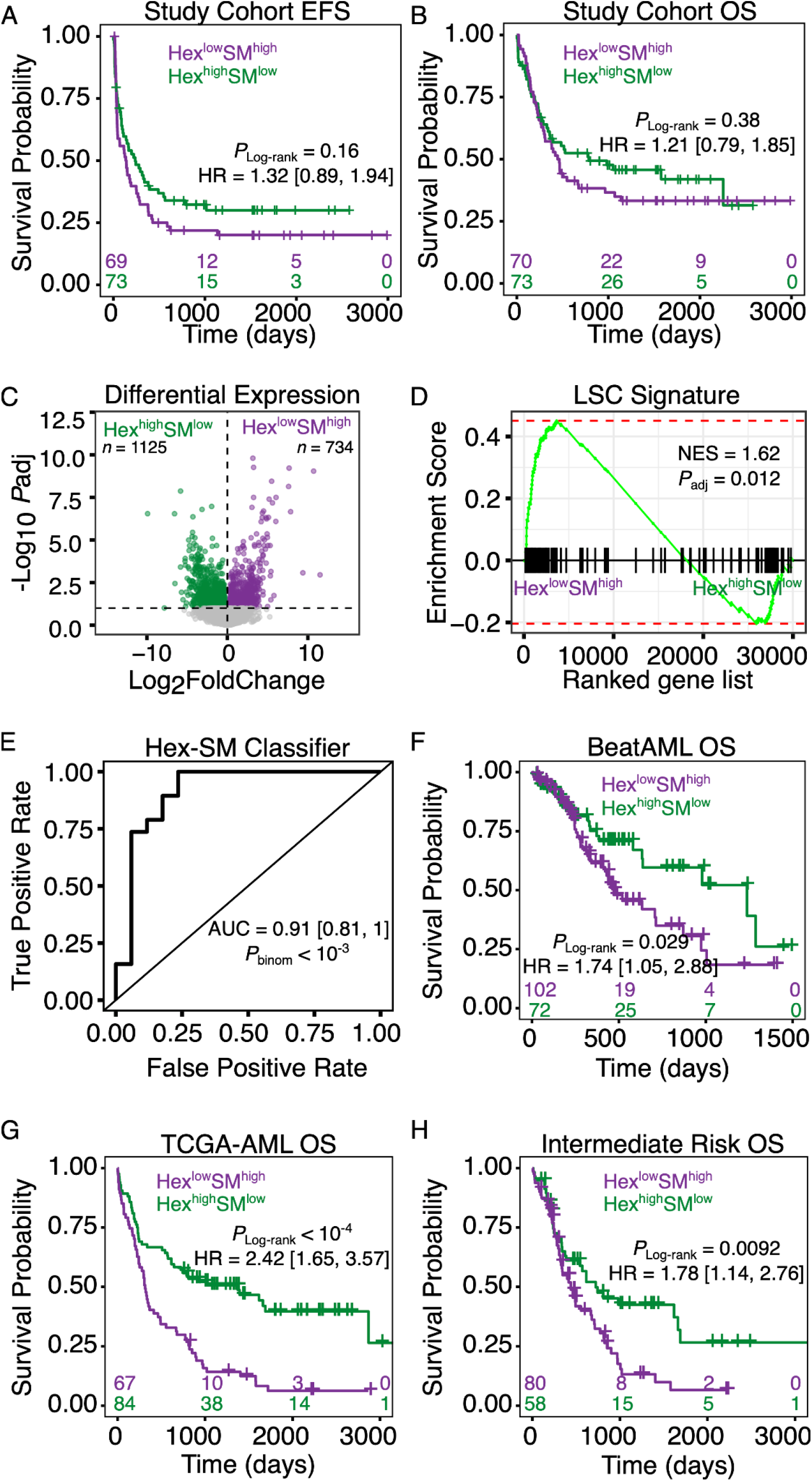
The Hex^low^SM^high^ and Hex^high^SM^low^ AML subtypes differ in gene expression and survival outcome. (**A-B**) Kaplan-Meier plots of event-free survival (EFS) (**A**) and overall survival (OS) (**B**) for AML patients grouped into Hex^low^SM^high^ (purple, *n* = 70) and Hex^high^SM^low^ (green, *n* = 73) subtypes. The study cohort comprised patients from three institutions (UVA, MSK, and PSU). Patients who received intensive induction chemotherapy treatment (“7+3”) were included in the analyses. Within each plot are the corresponding risk tables for the two groups. (**C**) Volcano plot of differentially expressed genes between the Hex^low^SM^high^ and Hex^high^SM^low^ subtypes. Purple genes are upregulated in the Hex^low^SM^high^ cluster whereas green genes are upregulated in the Hex^high^SM^low^ cluster. (**D**) The Hex^low^SM^high^ subtype is enriched for the leukemic stemness (LSC) program. Gene set enrichment analysis score plot for a leukemic stem cell signature of 104 genes^17^. The y-axis is the running enrichment score along the ranked gene list. The enrichment score is the maximum deviation from zero encountered in walking the list and represents the degree to which a gene set is over-represented at the top or the bottom of the ranked gene list. The normalized enrichment score (NES) is the enrichment score normalized for variation in gene set sizes. The adjusted *p*-value (*P*_*adj*_) for the NES is shown. (**E**) An RNASeq-based classifier accurately distinguishes sphingolipidomic subtypes. Receiver operating characteristics curve for a 284-gene support vector machine classifier applied to test data that includes both primary AML samples and cell lines with paired RNASeq and sphingolipidomic data. The area under the curve (AUC), its 95% confidence interval in brackets, and the one-sided binomial test *p*-value (*P*_binom_) of the classifier are shown. (**F-H**) Kaplan-Meier plots for AML patients inferred to be Hex^low^SM^high^ (purple) or Hex^high^SM^low^ (green) in BeatAML (*n*_purple_ = 102, *n*_green_ = 72) (**F**), TCGA-AML (*n*_purple_ = 67, *n*_green_ = 84) (**G**), and the molecularly defined Intermediate risk group combined for both BeatAML and TCGA-AML (*n*_purple_ = 80, *n*_green_ = 58) (**H**). Only AML patients who received standard intensive induction chemotherapy were included in the analyses for both datasets. Log-rank *p*-values, Hazard ratio (HR), and 95% confidence interval in brackets are shown. The bottom of each plot shows risk tables for the two subtypes.

To associate broader transcriptional differences with the subtypes, we collected RNASeq data for 29 primary AML samples and 30 AML cell lines with sphingolipidomic profiles. We appended RNASeq data from additional AML cell lines available through CCLE^14^, which were batch-corrected and merged with our data alongside two clinical RNASeq repositories for AML: TCGA-AML^15^ and BeatAML^16^ (**Supplemental Figure S2E-F**, see Methods). For the cell lines and primary samples with sphingolipidomics, we identified 734 transcripts increased in the Hex^low^SM^high^ subtype and 1125 transcripts increased in the Hex^high^SM^low^ subtype (**Figure 2C**, Supplemental Table 03; FDR-adjusted *P*_adj_ < 0.05), including five enzymes involved in sphingolipid metabolism (*UGCG, ST3GAL3, B3GALT1, FUT4, NAGA*). Genes characteristic of the clinically favorable Hex^high^SM^low^ subtype were enriched for hallmark gene sets related to immune activation (**Supplemental Figure S2G-H**). In contrast, genes for the clinically unfavorable Hex^low^SM^high^ subtype were enriched for four of five gene signatures associated with leukemic stem cells (LSCs)^17–20^ (**Figure 2D, Supplemental Figure S2I-L**). We concluded that the two sphingolipid subtypes were more coupled to transcriptomic states than AML driver mutations (**Supplemental Figure S1D**) and may relate to differences in biological mechanisms of the disease.

For inferring sphingolipidomic subtypes from transcriptional states alone, we developed a support vector machine classifier of Hex-SM status using the 284 most variable and differentially expressed genes between the subtypes (Supplemental Table 04). When trained on 60% of the samples with paired transcriptomics and sphingolipidomics (including both primary cases and AML cell lines), the classifier showed excellent predictive performance on the remaining 40% of samples (**Figure 2E**; area under the receiver operating characteristics curve = 0.91; balanced accuracy = 81%; one-sided binomial test *P* < 10^-3^). We then used the classifier with the batch-corrected transcriptomic data from TCGA-AML and BeatAML to infer sphingolipidomic subtypes. For both repositories, the classifier predicted a balanced proportion of Hex^low^SM^high^ and Hex^high^SM^low^ cases, supporting that neither subtype is rare. Consistent with our independent cohort (**Supplemental Figure S2B-C**), patients inferred to be Hex^low^SM^high^ had significantly worse survival outcomes than those predicted to be Hex^high^SM^low^ (**Figure 2F-G**; log-rank *P* = 0.029 for BeatAML, log-rank *P* < 10^-4^ for TCGA-AML). Among patients with molecularly defined risk classes in both datasets, we found that the Hex^high^SM^low^ subtype was enriched for the Favorable/Good group (Fisher’s exact test, *P* < 10^-4^), while the Hex^low^SM^high^ subtype was enriched for the Adverse/Poor group (Fisher’s exact test, *P* = 0.03; **Supplemental Figure S2M-N**). Interestingly, the Intermediate risk group was not detectably skewed by subtype (Fisher’s exact test, *P* = 0.07; **Supplemental Figure S2M-N**). Last, we stratified by risk group and examined whether patients in the two sphingolipid subtypes differ in their clinical outcomes. Overall survival of the Hex^low^SM^high^ subtype was similar to Hex^high^SM^low^ in the Favorable/Good risk group (log-rank *P* = 0.86), slightly worse in the Adverse/Poor risk group (log-rank *P* = 0.051), and significantly worse in the Intermediate risk group (log-rank *P* = 0.009; median OS = 439 vs. 723 days; **Supplemental Figure S2O-P; Figure 2H**). This extension to public AML datasets strengthens the conclusion that Hex^low^SM^high^ is a high-risk subtype with poor clinical outcomes, especially for patients whose molecular risk classification is Intermediate.

By examining AML from the perspective of sphingolipid metabolism, our work uncovers a stratification that eluded prior gene-based classifications^16,17,21^. The RNASeq-based classifier suggests that sphingolipidomic subtypes are embedded in a fraction of the transcriptome that is not prominent when clustering is performed in an unsupervised manner. Indeed, the sphingolipid profiles of AML cell lines are much more concordant with primary samples (**Figure 1C**), unlike when their whole transcriptomes are co-clustered (**Supplemental Figure S3A-C**). The stemness transcriptional programs enriched in the high-risk Hex^low^SM^high^ subtype are consistent with the importance of sphingolipid homeostasis in maintaining hematopoietic stem cells^13^. Since gene expression panels are used clinically^22^, our sphingolipid-guided transcript classifier could be useful for identifying patients least likely to benefit from the intensive induction combination chemotherapy and thus most eligible to receive experimental therapeutics. Future studies should confirm these findings and investigate the pharmacologic vulnerabilities of the two sphingolipid subtypes. Given the promise of targeting sphingolipid metabolism in AML^7,23,24^, we envision that sphingolipidomic subtyping could contribute to tailored treatment selections for AML patients that otherwise lack targetable alternatives.

## Acknowledgments

We thank Samuel Haddox for help with RNA preparation for RNASeq, Emily Sullins for processing patient and cell lines, Galina Diakova for help with RNASeq data from The Oncology Research Information Exchange Network, and Memorial Sloan Kettering Tissue Banks for assistance with sample processing. We appreciate comments from Dr. Cameron Griffiths, Wisam Fares, and Russell Hawes on the manuscript figures.

This work was supported by the National Institutes of Health (NIH) under the National Cancer Institute (NCI) Award Number P01 CA171983 (to TPL and MK), NIH/NCI Cancer Center Support Grant P30 CA044579 (to TPL), NCI R35 CA197594 (to RLL), NIH/NCI Cancer Center Support Grant P30 CA008748 (to RLL), NIH/NCI F31 CA271809 and UVA Robert R. Wagner Fellowship (to JU), NCI K08 CA215317 and Edward P. Evans Foundation (to ADV), and NCI R03 CA252825 (BMB).

## Authorship

## Contribution

- designed research: BBP, SFT, TEF, MC, FGB, MK, DC, DJF, KAJ, TPL
- performed research: BBP, SFT, TEF, WD, IL, AS, AV, DC
- contributed vital new reagents or analytical tools: TEF, FGB
- collected data: SFT, TEF, JU, WD, IL, AS, AV, FGB, DC, DJF
- analyzed and interpreted data: BBP, SFT, TEF, JU, JS, BMB, MST, MC, FGB, RLL, DC, DJF, KAJ, TPL
- performed statistical analysis: BBP, FGB, KAJ
- wrote the manuscript: BBP, SFT, TEF, JU, FGB, DJF, KAJ, TPL. All authors read and approved the manuscript.

## Figure Legends for Supplementary Figures

**Supplemental Figure S1:**
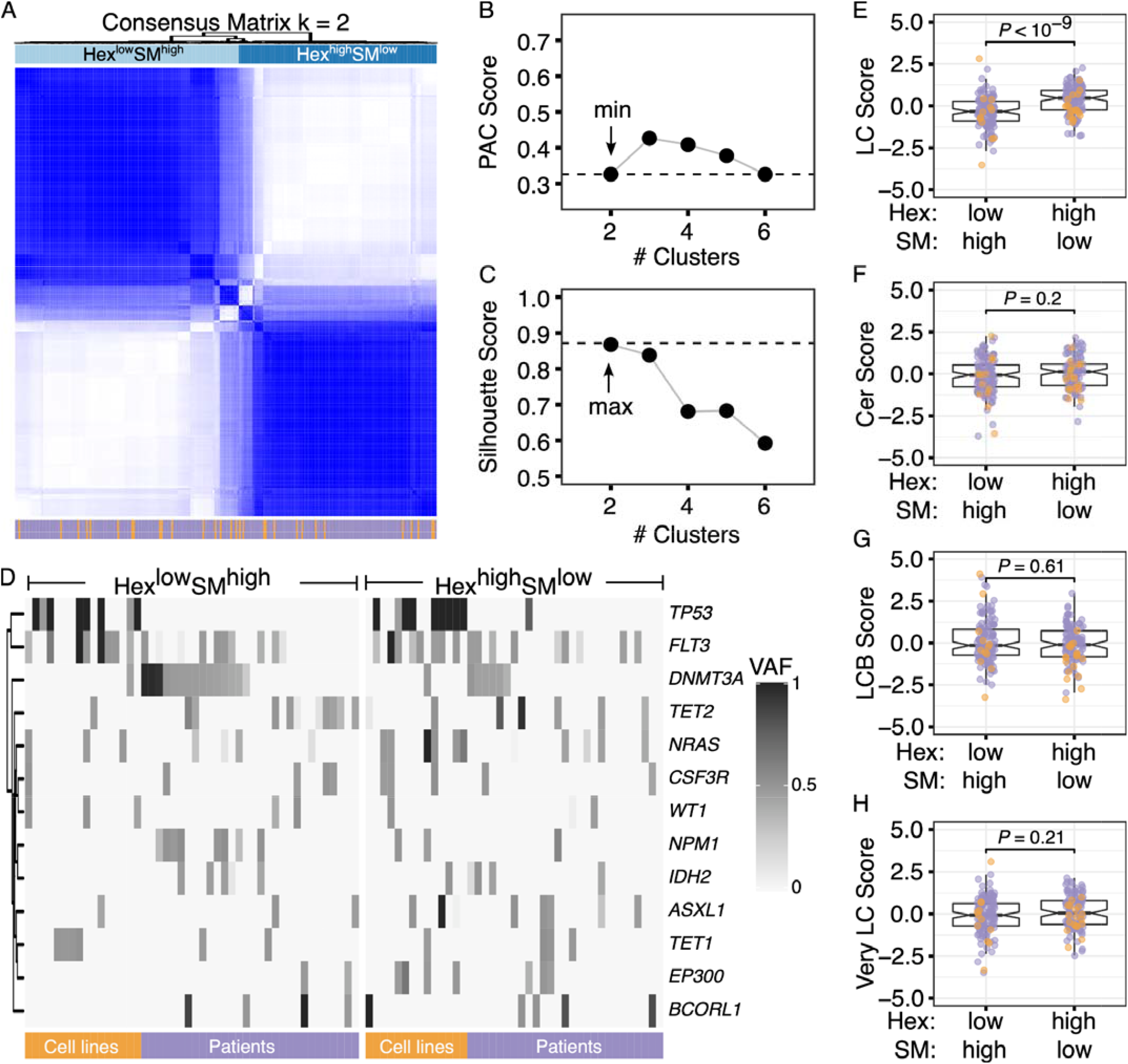
Genomic and sphingolipidomic associations within the two AML consensus clusters. (A) Consensus clustering on normalized lipidomics data was used to identify stable sphingolipid clusters in human AML primary samples and AML cell lines. The consensus score heatmap for two clusters (*k* = 2) is shown, illustrating the frequency that a given pair of samples was placed in the same cluster over 1,000 iterations. Each point denotes a pair of samples colored by consensus score from white (0, never co-cluster) to blue (1, always co-cluster). Colors denote primary AML samples (purple, *n* = 213) or AML cell lines (orange, *n* = 30). (**B-C**) Cluster statistics supporting two stable clusters. Proportions of ambiguous clustering (PAC) scores (**B**) and Silhouette scores (**C**) for clusters *k* = 2 to *k* = 6. The cluster with the lowest PAC and highest Silhouette score was chosen as the optimum sphingolipid cluster for AML samples. (**D**) Two sphingolipid metabolic clusters do not differ in their mutational profiles. Heatmap for the estimated variant allele frequency (VAF) for genes detected as mutated (VAF > 0) in over 10% of AML samples. Samples (columns) are separated based on their sphingolipid cluster, either Hex^low^SM^high^ or Hex^hgh^SM^low^; colors indicate primary AML samples (purple, *n* = 57), and AML cell lines (orange, *n* = 30). No differences in mutation frequency were detected between the clusters by Fisher’s exact test (*P≥*0.27). (**E-H**) Normalized Z-scores for lipid species within the long-chain (C14-C20-carbon) sphingolipid species (**E**), Ceramide (Cer) (**F**), sphingolipid long-chain/sphingoid bases (LCB) (**G**), or very-long-chain (C22-C26-carbon) sphingolipid species (**H**) were summed and differences between the two sphingolipid consensus clusters were assessed by Mann-Whitney test with continuity correction. Colors indicate primary AML samples (purple, *n* = 213), and AML cell lines (orange, *n* = 30).

**Supplemental Figure S2:**
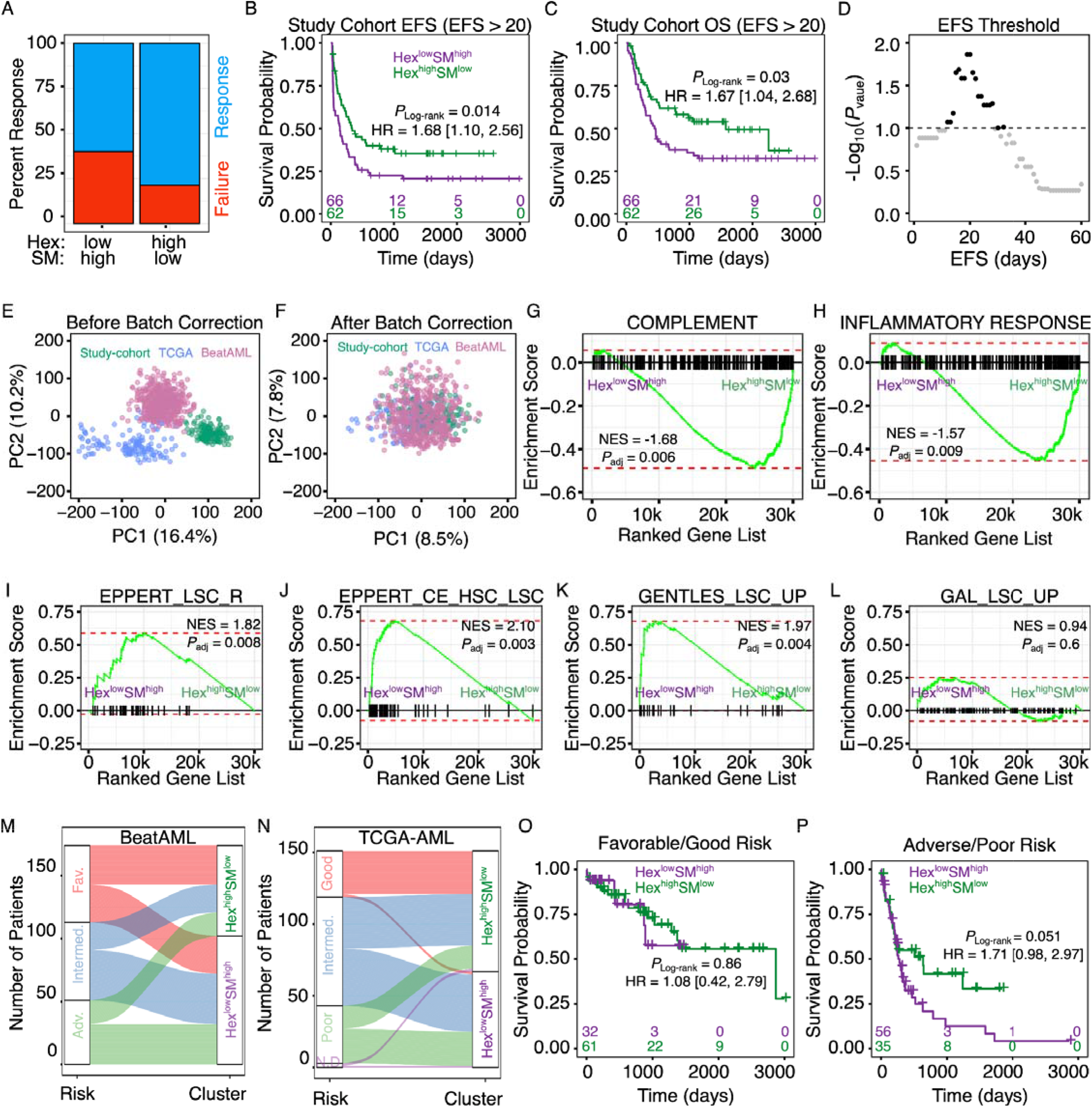
Gene expression and survival outcome differences between the two sphingolipidomic AML subtypes. (**A**) Barplot for proportions of AML patients grouped into Hex^low^SM^high^ (*n* = 65) and Hex^high^SM^low^ (*n* = 66) subtypes with response or failure to intensive induction combination therapy. The difference in response between the clusters was significant by Fisher’s exact test (*P* = 0.02). (**B-C**) Kaplan-Meier plots for AML patients grouped into Hex^low^SM^high^ (purple) and Hex^high^SM^low^ (green) subtypes for patients in the study cohort with the exclusion of early death or events within the first 20 days of intensive induction chemotherapy treatment; event-free survival (**B**), overall survival (**C**). Log-rank *p*-values, Hazard ratio (HR), and 95% confidence interval in brackets are shown. The bottom of each plot shows risk tables for the two subtypes. (**D**) Sensitivity of the EFS threshold for discriminating differences between the two clusters in EFS for the study cohort. The -log_10_ of log-rank *p*-values is plotted versus the minimum EFS day for thresholding. The dotted line denotes *P*_Log-rank_ = 0.1. (**E-F**) Integration of RNASeq data from the study cohort, TCGA, and BeatAML. Principal component analysis (PCA) with the top 50% of most variable genes before (**E**) and after (**F**) batch correction. The data were normalized using DESeq2 and log-transformed with log_2_ (normalized counts + 1). Colors indicate the source of the data, and study cohort (green, *n* = 148), TCGA (blue, *n* = 151), and BeatAML (pink, *n* =510). (**G-H**) Hallmark complement pathways (**G**), and Hallmark inflammatory response (**H**) are enriched in the trailing genes upregulated in Hex^high^SM^low^. (**I-L**) The Hex^low^SM^high^ subtype is enriched for the leukemic stem cell (LSC) program. Gene set enrichment score plot for genes up-regulated in functionally defined LSC from AML patients^18^ (**I**), genes shared between hematopoietic stem cells (HSC) and AML LSC genes^18^ (**J**), genes up-regulated in LSC compared to leukemic progenitor cells from AML patients^20^ (**K**), genes upregulated in leukemic stem CD34+CD38-cells from AML compared to the CD34+CD38+ cells^19^ (**L**). The y-axis is the running enrichment score (ES) along the ranked gene list. The enrichment score is the maximum deviation from zero encountered in walking the list and represents the degree to which a gene set is over-represented at the top or the bottom of the ranked gene list. The normalized enrichment score (NES) is the ES normalized for variation in gene set sizes. The adjusted *p*-value (*P*_*adj*_) for the NES is shown. (**M-N**) Proportional distributions of molecularly defined risk classification in the two sphingolipidomic subtypes for BeatAML with European LeukemiaNet (ELN) 2017 categories (**M**), and TCGA-AML with molecularly defined risk categories (**N**). (**O-P**) Kaplan-Meier plots for overall survival of AML patients predicted to be Hex^low^SM^high^ (purple) and Hex^high^SM^low^ (green) subtypes for patients in the BeatAML, and TCGA-AML separated by their molecularly defined risk classifications: Favorable/Good risk (**O**) and Adverse/Poor risk (**P**). Log-rank *p*-values, Hazard ratio (HR), and 95% confidence interval in brackets are shown. The bottom of each plot shows risk tables for the two subtypes.

**Supplemental Figure S3:**
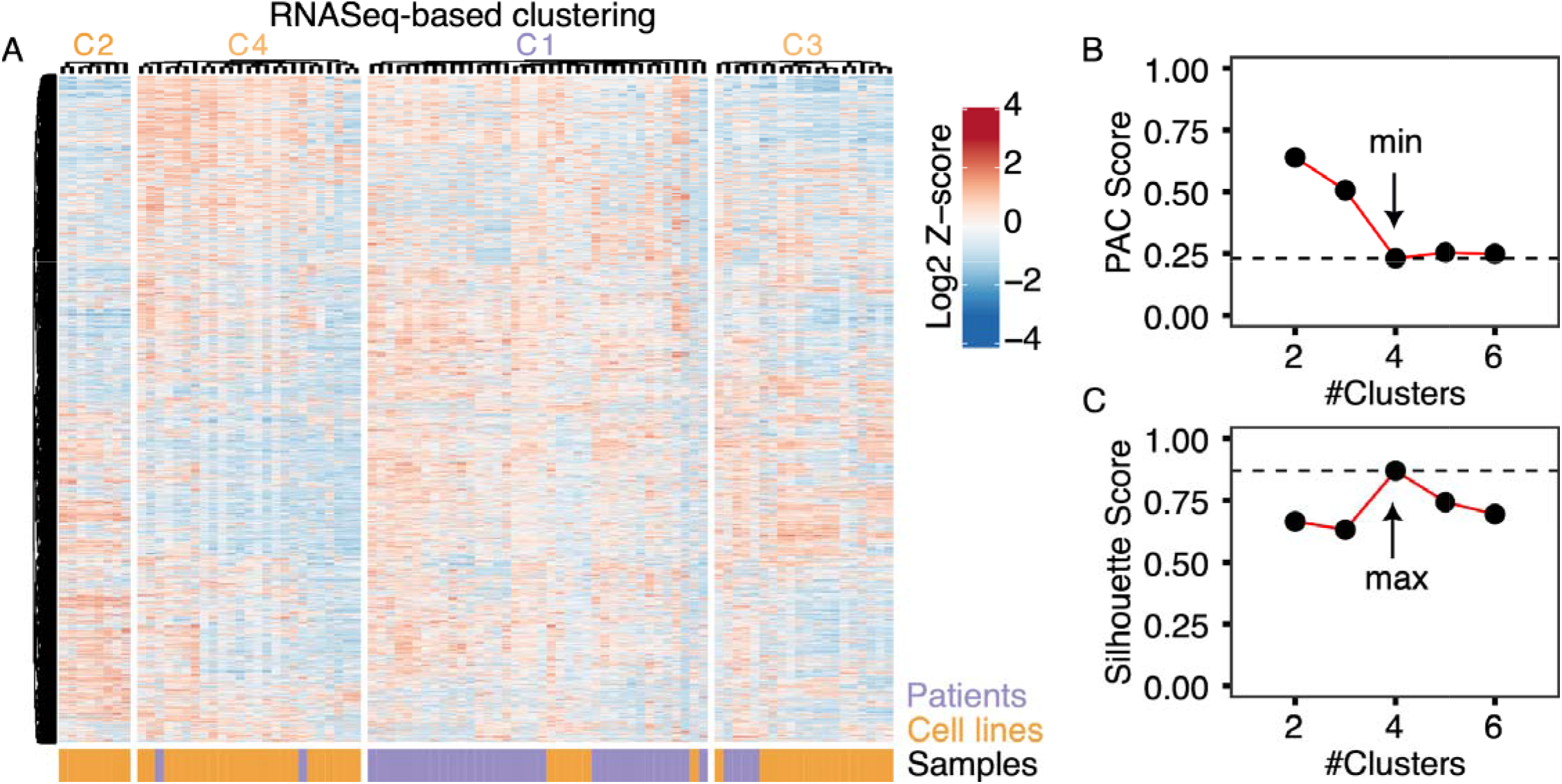
Whole transcriptomes separate AML samples into four stable clusters that segregate cell lines and patient samples. (**A**) Sample-to-sample differences in row standardized expression of 2000 most variably expressed genes, separated by consensus clusters based on transcriptomic data of AML cell lines and primary samples. Expression values are from batch-adjusted, DESeq2-normalized data and log-transformed with log_2_ (normalized counts + 1). Colors indicate the sample type: AML cell lines (orange, *n* = 53) and primary samples (purple, *n* = 38). C1 is enriched in patient samples (*P <* 10^-11^), whereas C2, C3 and C4 are enriched in cell lines (C2: *P* = 0.01; C3: *P* = 0.03; C4: *P* < 10^-4^ ; Fisher’s exact test). (**B-C**) Cluster statistics supporting four stable clusters based on RNASeq data. Proportions of ambiguous clustering (PAC) scores (**B**) Silhouette scores (**C**) for clusters, *k* = 2 to *k* = 6. The cluster with the lowest PAC and highest Silhouette score was chosen as the optimum sphingolipid cluster for AML cell lines and primary samples.

