## Supplementary Methods for "Acute myeloid leukemia stratifies as two clinically relevant sphingolipidomic subtypes"

^1^Department of Biomedical Engineering, University of Virginia, Charlottesville, VA. ^2^Department of Medicine, Division of Hematology & Oncology, University of Virginia, Charlottesville, VA. ^3^Department of Pharmacology, University of Virginia, Charlottesville, VA. ^4^Department of Microbiology, Immunology & Cancer Biology, University of Virginia, Charlottesville, VA. ^5^Department of Pharmacology, Pennsylvania State University College of Medicine, Hershey, PA. ^6^Penn State Cancer Institute, Hershey, PA. ^7^Departments of Medicine, Division of Hematology & Oncology, and of Genetics & Development, Columbia Stem Cell Initiative, Herbert Irving Comprehensive Cancer Center, Columbia University Irving Medical Center, New York, NY. ^8^Department of Chemistry, Biology, and Health Sciences, South Dakota School of Mines and Technology, Rapid City, SD. ^9^Northwestern University Feinberg School of Medicine Robert H. Lurie Comprehensive Cancer Center Chicago, IL. ^10^Department of Biochemistry & Molecular Biology, Brody School of Medicine, and the East Carolina Diabetes and Obesity Institute, East Carolina University, Greenville, NC. ^11^Department of Biochemistry and Molecular Genetics, University of Virginia, Charlottesville, VA. ^12^University of Virginia Comprehensive Cancer Center, Charlottesville, VA. ^13^Human Oncology and Pathogenesis Program and Leukemia Service, Department of Medicine, Memorial Sloan Kettering Cancer Center, New York, NY. ^14^Department of Medicine, Division of Hematology and Oncology, Pennsylvania State University College of Medicine, Hershey, PA. (*deceased; # corresponding authors)

**Corresponding Authors (#):**

Thomas P. Loughran, Jr., M.D.

Department of Medicine, Division of Hematology & Oncology,

The University of Virginia School of Medicine, Charlottesville, VA, 22908

Kevin A. Janes, Ph.D.

Department of Biomedical Engineering, Department of Biochemistry & Molecular Genetics

The University of Virginia, Charlottesville, VA, 22908

**SUPPLEMENTAL FIGURES AND TABLES:**

[Supplementary Tables](https://docs.google.com/spreadsheets/d/1HOjBkH1eKEndszdOO1ojUo50ITEE7pdF/edit#gid=1216893570)

Supplementary Figures

**MATERIALS AND METHODS:**

***Sample Acquisition and Handling:*** Samples from AML patients without prior treatment were obtained by informed consent for sample collection according to the Declaration of Helsinki and protocols approved by the Institutional Review Boards of Penn State Hershey Cancer Center, Memorial Sloan Kettering Cancer Center, and the University of Virginia Cancer Center (Oncology Research Information Exchange Network [ORIEN] program). Normal bone marrow samples were purchased commercially (All Cells, LLC, Emeryville, CA). All blood or bone marrow samples were enriched for mononuclear cells with Ficoll-Pague PLUS (Cytiva, Marlborough, MA) using the gradient separation method^1^.

***Lipid Extraction and Sphingolipid Profiling:*** Human AML cell lines were cultured as previously described^2^. Cells were seeded at 7.5x10^5^ cells/mL, cultured for 24 hours, and pelleted/washed in 1x PBS before lipid extraction. Frozen cryopreserved primary AML patient samples were thawed and washed with 1x PBS, followed by lipid extraction from cell homogenates using a mix of isopropanol:water:ethyl acetate (3:1:6; v:v:v). Internal standards (10 pmol of d17 long-chain bases and C12 acylated sphingolipids) were added to samples at the onset of the extraction procedure. Extracts were separated on a Waters I-class Acquity UPLC chromatography system. Mobile phases were (A) 60:40 water:acetonitrile and (B) 90:10 isopropanol:methanol with both mobile phases containing 5 mM ammonium formate and 0.1% formic acid. A Waters C18 CSH 2.1 mm ID × 10 cm column maintained at 65°C was used to separate the sphingoid bases, 1-phosphates, and acylated sphingolipids. The eluate was analyzed with an inline Waters TQ-S mass spectrometer using multiple reaction monitoring. Lipidomics data for each run was log_2_-transformed and mean-centered across samples for each sphingolipid species. The mean-centered log_2_-transformed data from different runs were combined to create the final normalized lipidomics dataset.

***Variant Allele Frequency Profiling:*** Human AML cell lines and patient samples were sequenced using targeted (585 genes) high-throughput next-generation sequencing (MSK-IMPACT Heme) as previously described^3^. Mutation calling was performed to identify single-point variants using Mutect and calling of insertions and deletions using Pindel as described previously^3^. The sequences were aligned to pooled normal bone marrow samples and then filtered for common variants and single nucleotide polymorphisms. Mutations were excluded if present in at least one database of known non-somatic variants (dbSNP and 1000 genomes) and absent from COSMIC. Non-excluded mutations with variant allele frequency > 0.02 were called as a mutation.

***Consensus Clustering and Statistics:*** Consensus clustering^4^ was performed on normalized lipidomics data or transcriptomics data for AML cell lines and primary samples using ConsensusClusterPlus (version 1.52.0) in R with 80% sub-sampling of lipid samples, 1000 repetitions, 1 - Pearson correlation, and the partition around medoids clustering algorithm with innerLinkage and finalLinkage both set to complete. Proportions of Ambiguous Clustering (PAC) score was calculated using the PAC function of diceR (version 1.2.2) in R with the Consensus Matrix from ConsensusClustering, where lower and upper bounds were set to 0.1 and 0.9 respectively. Silhouette scores were calculated for clustering results from *k* = {2, 10} using the silhouette_SimilarityMatrix function of CancerSubtypes (version 1.14.0) in R. The cluster with the highest Silhouette score and the lowest PAC score was selected as the optimum cluster for further analysis.

***Lipid Class Definition:*** The identified consensus clusters were defined by the abundance of lipid species. Briefly, lipid species were categorized into four main classes: Hexosylceramides (Hex, comprised of glucosyl- and galactosylceramides), Sphingomyelin (SM), Ceramide (Cer), and Long-Chain Bases (LCB) (comprised of sphingosine, dihydrosphingosine, sphingosine-1-phosphate, hexosylsphingosine). Normalized lipid Z-scores within each lipid class were added to calculate Hex, SM, Cer, and LCB scores respectively for each AML sample. Similarly, sphingolipid species were grouped into two categories based on carbon chain length (C14-C20, long-chain lipids; C22-C26, very long-chain lipids), and normalized Z-scores within each lipid chain class were added to calculate either long-chain or very long-chain lipid scores for each AML sample. Differences between the clusters were assessed by the Mann-Whitney test with continuity correction.

***RNA-Seq Data:***

***CCLE-RNASeq data****:* RNA-Seq data for human AML cell lines (Supplemental Table 05, Column #1) from Cancer Cell Line Encyclopedia were downloaded as BAM files from the Genomic Data Commons (GDC) data portal of the National Cancer Institute (NCI) using the GDC-Client tool. The corresponding BAM files were converted to fastq files using samtools (version 1.12, using htslib 1.12) with the default parameters.

***UVA-RNASeq data1***: RNA-Seq for human AML cell lines (Supplemental Table 05, Column #2) was performed by Psomagen. Briefly, 100 ng RNA isolated by trizol standard protocol (according to the manufacturer’s instructions) was used for sequencing with TruSeq Stranded Total RNA Ribo Zero kit (Human/Mouse/Rat). Samples were sequenced as 150 bp paired-end reads on a NovaSeq 6000 S4 with a sequencing depth between 20–39 million reads per sample.

***UVA-RNASeq data2****:* RNA-Seq for human AML cell lines and primary samples (Supplemental Table 05, Column #3) was performed by Novogene Co., Ltd. Briefly, total RNA was isolated from AML patient PBMCs and AML cell lines by resuspending pellets in Trizol and purifying RNA with Direct-zol RNA Miniprep (Zymo Research, Irvine, CA). DNase treatment was done using the RNase-Free DNase Set (Qiagen # 79254). RNA integrity was analyzed on an Agilent TapeStation system according to the manufacturer’s protocol. RNA samples with RIN scores above 5.3 were sent to Novogene for mRNA library prep and sequencing. Briefly, messenger RNA was purified from total RNA using poly-T oligo-attached magnetic beads. After fragmentation, the first strand cDNA was synthesized using random hexamer primers followed by the second strand cDNA synthesis. The library was sequenced after end repair, A-tailing, adapter ligation, size selection, amplification, and purification. Samples were sequenced as 150 bp paired-end reads on a NovaSeq 6000 S4 with a sequencing depth between 20– 57 million reads per sample.

***ORIEN-RNASeq data****:* RNA-Seq data for human AML primary samples (Supplemental Table 05, Column #4) were downloaded from ORIEN as cram files. The downloaded cram files were converted to bam files and then to fastq files using samtools (version 1.12, htslib 1.12) with the default parameters.

***TCGA-AML RNASeq data****:* RNA-Seq data for human AML primary samples from the TCGA-AML cohort were downloaded from NCI’s GDC data portal as htseq read counts. Patient IDs for the TCGA cohort used in this study are listed in Supplemental Table 05, Column #5.

**BeatAML RNASeq data**: RNA-Seq data for human AML primary samples from BeatAML 1.0-Cohort was downloaded from NCI’s GDC data portal as htseq read counts. Patient IDs for the BeatAML used in this study are listed in Supplemental Table 05, Column #6.

***RNA-Seq Processing and Analysis***: For fastq files, read quality was assessed with FastQC (version 0.11.5), and Illumina adapter sequences were trimmed using Trimmomatic (version 0.39). The trimmed fastq files were aligned to the GRCh38 human reference genome (hg38, release 86) using HISAT2 (version 2.1.0)^5^ and gene counts were obtained using featureCounts, downloaded from Subread^6^. All downstream analyses were performed in R Studio (version, 1.4.1106) using the Bioconductor framework. To integrate RNASeq from different sources, only transcripts for which counts were available in all datasets were retained for further analyses. Read counts were batch adjusted with the Combat_seq function of sva (version 3.36.0) in R using default parameters. The final batch-adjusted data contained 56,485 transcripts that were present in all processed read counts.

***Differential Gene Expression Analysis:*** Differential gene expression analyses between sphingolipid clusters (Hex^low^SM^high^ vs. Hex^high^SM^low^) were performed using the DESeq2 package (version 1.28.1) in R with batch-adjusted count data on AML cell lines and primary samples for which the complementary lipidomics data were available. The adjusted count data were normalized using the ‘counts’ function in DESeq2. All downstream analyses were performed with DESeq2-normalized data.

***Lipid-Gene Classifier:*** To select the most informative genes for the classifier, the normalized count was reduced to the top 50% of most variable genes and quantified by median absolute deviation by using the R function ‘mad’. Gene signatures were further reduced to include only genes that were differentially expressed (absolute log_2_ fold change > 0 & adjusted *P*-value < 0.01) between two sphingolipid metabolic clusters. The final classifier consisting of 284 genes was built as a support vector machine (SVM) with linear kernel implementation in the R package caret (version 6.0-93) using the default parameters. The SVM classifier was trained on 60% of random samples (consisting of both cell lines and primary samples) with both omics data types. The model was tested on 40% of testing data containing both cell lines and primary samples for which lipidomics and RNASeq were available. The accuracy of the classifier was assessed by ROC curve, Sensitivity, Specificity, and Balanced accuracy. The model had a balanced accuracy of 0.81, with a confidence interval [0.64, 0.92], and *p*-value of 0.00052, Sensitivity: 0.82 & Specificity: 0.80, the area under the ROC curve = 0.90 [0.82, 0.99].

***Pathway and Gene Set Enrichment Analysis:*** The package mgsigdr (version 7.5.1) was used in R to retrieve Hallmark terms from the Human Molecular Signature Database (MSigDB) Collections, and the package fgsea (version 1.14.0) was used to perform gene set enrichment analysis on ranked gene lists with arguments, minSize = 2, maxSize = 1000, nPerm = 10000, and other arguments left at the default settings. All genes from differential expression analyses with baseMean ≥ 1 were included and ranked by their log2 fold change. Pathways were considered enriched if adjusted P-values < 0.05. Leukemic stem cell (LSC) signatures consisted of 104 differentially expressed genes^7^, genes shared between hematopoietic stem cell (HSC) and AML LSC genes^8^, genes up-regulated in functionally defined LSC from AML patients^8^, genes up-regulated in LSC compared to leukemic progenitor cells from AML patients^9^, and genes up-regulated in LSC defined as CD34+CD38- cells from AML patients compared to CD34+CD38+ cells^10^.

***Survival Analysis:*** Survival analyses were performed with the survfit and coxph functions of the package survival (version 3.4-0) in R. Survival plots were generated using the function ggsurvplot of the package survminer (version 0.4.9) in R.

***Data Visualization:*** Heatmaps were generated using the Heatmap function of the package ComplexHeatmap (version 2.4.3) in R using color palettes “Greys” and “RdBu” with the brewer.pal function of the Package RColorBrewer (version 1.1-3) in R. The package ggplot2 (version 3.3.6) was used for data visualization unless stated otherwise. The UMAP was generated with the umap function of the package uwot (version 0.1.14) in R.

***Data Availability:*** RNASeq data is available at the Gene Expression Omnibus (GSE229032; <https://www.ncbi.nlm.nih.gov/geo/query/acc.cgi?acc=GSE229032> reviewer token: mhmjocsmfzcvnwx). Normalized lipidomics data is provided in Supplemental Information (Supplemental Table 06). Some data on primary AML samples analyzed in this study (identifiers in Supplemental Table 05, Column #4) are subject to restrictions set by the ORIEN network, and access is controlled by M2Gen and the ORIEN consortium. Requests to access these datasets should be directed to [https://www.oriencancer.org/request-an-accoun](https://www.oriencancer.org/request-an-account)t.

***Statistics:*** Statistical tests are reported in the figure legends or text as appropriate.
